# Effect of sodium silicate on drinking water biofilm development

**DOI:** 10.1101/2021.09.15.460503

**Authors:** Sebastian Munoz, Benjamin F. Trueman, Bofu Li, Graham A. Gagnon

## Abstract

Sodium silicates have been studied for sequestration of iron, coagulation, and corrosion control, but their impact on biofilm formation has not been documented in detail. This study investigated the impact of sodium silicate corrosion control on biomass accumulation in drinking water systems in comparison to orthophosphate, a common corrosion inhibitor. Biofilm growth was measured by determining ATP concentrations, and the bacterial community was characterized using 16S ribosomal RNA (rRNA) sequencing. A pilot-scale study with cast-iron pipe loops, annular reactors (ARs), and polycarbonate coupons demonstrated significantly lower biofilm ATP concentrations in the sodium silicate-treated AR than the orthophosphate-treated AR when the water temperature exceeded 20°C. However, an elevated sodium silicate dose (48 mg L^-1^ of SiO_2_) disturbed and dispersed the biofilm formed inside the AR, resulting in elevated effluent ATP concentrations. Two separate experiments confirmed that biomass accumulation was higher in the presence of orthophosphate at high water temperatures (20°C) only. No significant differences were identified in biofilm ATP concentrations at lower water temperatures (below 20°C). Differences in bacterial communities between the orthophosphate- and sodium silicate-treated systems were not statistically significant, even though orthophosphate promoted higher biofilm growth. However, the genera *Halomonas* and *Mycobacterium*—which include opportunistic pathogens—were present at greater relative abundances in the orthophosphate-treated system compared to the sodium silicate system.

**Graphical abstract:** 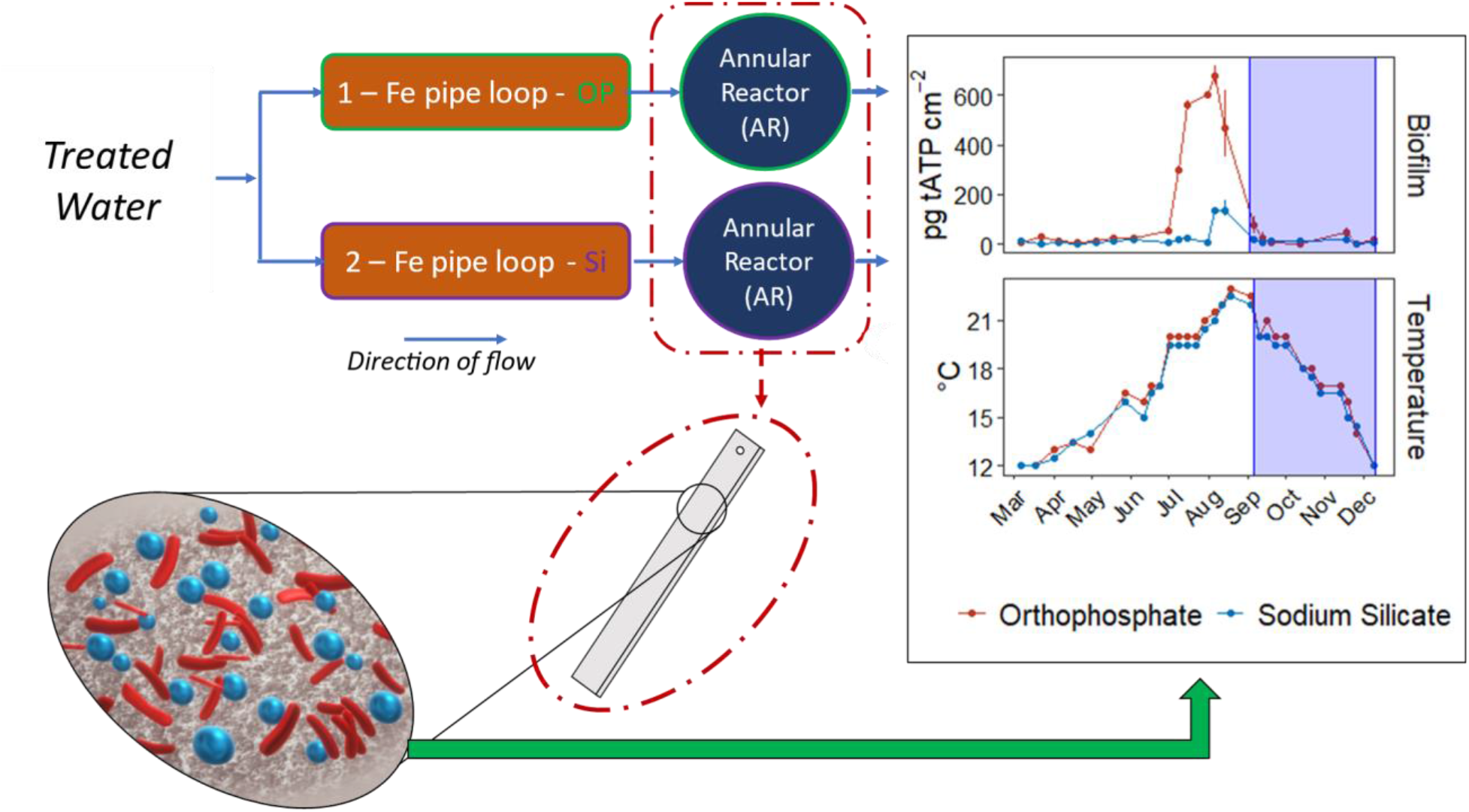

Orthophosphate promotes more biofilm growth in comparison to sodium silicates at water temperatures above 20°C.

**Water impact statement:** Sodium silicates have been used in drinking water treatment for decades, both as sequestrants and as corrosion inhibitors. However, their impact on biofilm formation is poorly understood, and this risks drinking water quality. This study aims to further clarify the effects of corrosion inhibitors on biofilm development, including inhibitors that are not phosphate-based.

## Introduction

Internal corrosion of drinking water distribution pipes can release lead, copper, and iron to drinking water. It can also result in tubercles—deposited corrosion products—which accumulate biofilm and provide an environment for bacteria and opportunistic pathogens.^1,2^ Corrosion by-products can react with disinfectant, neutralizing it before it can inactivate microorganisms in the bulk water or biofilm.^1,3,4^ To mitigate corrosion, drinking water treatment plants apply corrosion inhibitors while making adjustments to pH and alkalinity. According to a 2019 corrosion control survey, 54% of utilities in the US utilize a phosphate-based corrosion inhibitor.^5^

While in most cases organic carbon limits biofilm growth in drinking water distribution systems, phosphorus has also been identified as the growth-limiting nutrient, promoting biofilm regrowth.^6–9^ Phosphate may promote bacterial diversity as well as biofilm growth,^7,10^ but some work contradicts this finding.^11^ Other studies suggest that phosphate neither influences total bacterial densities,^2,12^ nor supports biofilm growth.^13^ These inconsistencies motivate further investigation of the effects of corrosion inhibitors on biofilm development, including inhibitors that are not phosphate-based.

Besides orthophosphate, some water utilities use sodium silicates as corrosion inhibitors. While sodium silicates are mostly used for sequestration of iron and coagulation applications, they have been studied for lead release control applications.^14– 16^ Sodium silicates may also inhibit the oxidation of ferrous iron, particularly in the pH range of 6.0 – 7.0,^17^ and they may reduce color and water turbidity.^18^

While some focus has been directed to understanding sodium silicates as corrosion inhibitors, their impact on biofilm formation has not been documented in detail. A few studies do suggest that silicate treatment exerts a minimal influence on biofilm: Rompré *et al*.^2^ reported that sodium silicate did not influence biofilm growth in comparison to orthophosphate and blended ortho-polyphosphate inhibitors, suggesting that biofilm accumulation is influenced more by the substrate material than the corrosion inhibitor. Similarly, Kogo *et al*.^19^ reported no significant difference in adenosine triphosphate (ATP) concentrations from biofilm samples among the orthophosphate, zinc orthophosphate, and sodium silicate-treated systems.

The main objective of this work was to compare the impact of sodium silicate on biomass accumulation in drinking water systems against the frequently used orthophosphate, as measured by ATP concentrations and 16S ribosomal RNA (rRNA) sequencing analysis for bacterial community characterization.

A pilot-scale study is presented in this paper to describe the impacts of sodium silicates on biofilm growth using water from a drinking water treatment plant. Our objectives were:

1. To study biofilm growth in the presence of sodium silicate, using a pilot-scale model distribution system comprising cast-iron pipe loops feeding rotating annular reactors (ARs). Biomass accumulation inside the ARs was monitored over a 10-month period that included seasonal water temperature variation.
2. To monitor, in the same model system, the biomass accumulated at low water temperatures (below 20°C) on polycarbonate coupons representing PVC and cast-iron pipe loops treated with sodium silicate or orthophosphate.

A bench-scale experiment with the same filtered water used in the pipe loop system was completed using batch reactors and polyethylene (PEX) coupons. The objectives of this experiment were:

1. To compliment the findings from the pilot study by measuring the biomass accumulated on PEX coupons treated with sodium silicate or orthophosphate.
2. To replicate the biomass accumulation on coupons exposed to high water temperatures (20°C).

## Materials and methods

### Model distribution system

#### Pipe loop system

Filtered water from J.D. Kline Water Supply Plant (JDKWSP) (Halifax, Nova Scotia, Canada) was used as feed water for the pilot distribution system (the feedwater contained no orthophosphate). JDKWSP uses alum coagulation followed by dual media direct filtration (sand and anthracite). A detailed description of the facility can be found in Knowles *et al*.^20^, and Stoddart *et al*.^21^. The model distribution system comprised four independent pipe loops: two unlined cast-iron and two PVC pipe distribution main sections arranged in parallel (Figure 1) and designed to simulate water aging in a drinking water distribution system. The iron distribution main sections were 150 mm in diameter and moderately to heavily tuberculated, whereas the PVC sections were 100 mm in diameter. Both types of distribution mains were approximately 1.8 m in length. The hydraulic retention time (HRT) in the loops was set at 12 hours, with water flowing in through the main pipes at a rate of 0.03 m s^-1^ (approximate values). A detailed description of the pipe loop apparatus can be found in Gagnon *et al*.^22^, and Woszczynski *et al*.^23^.

**Figure 1:**
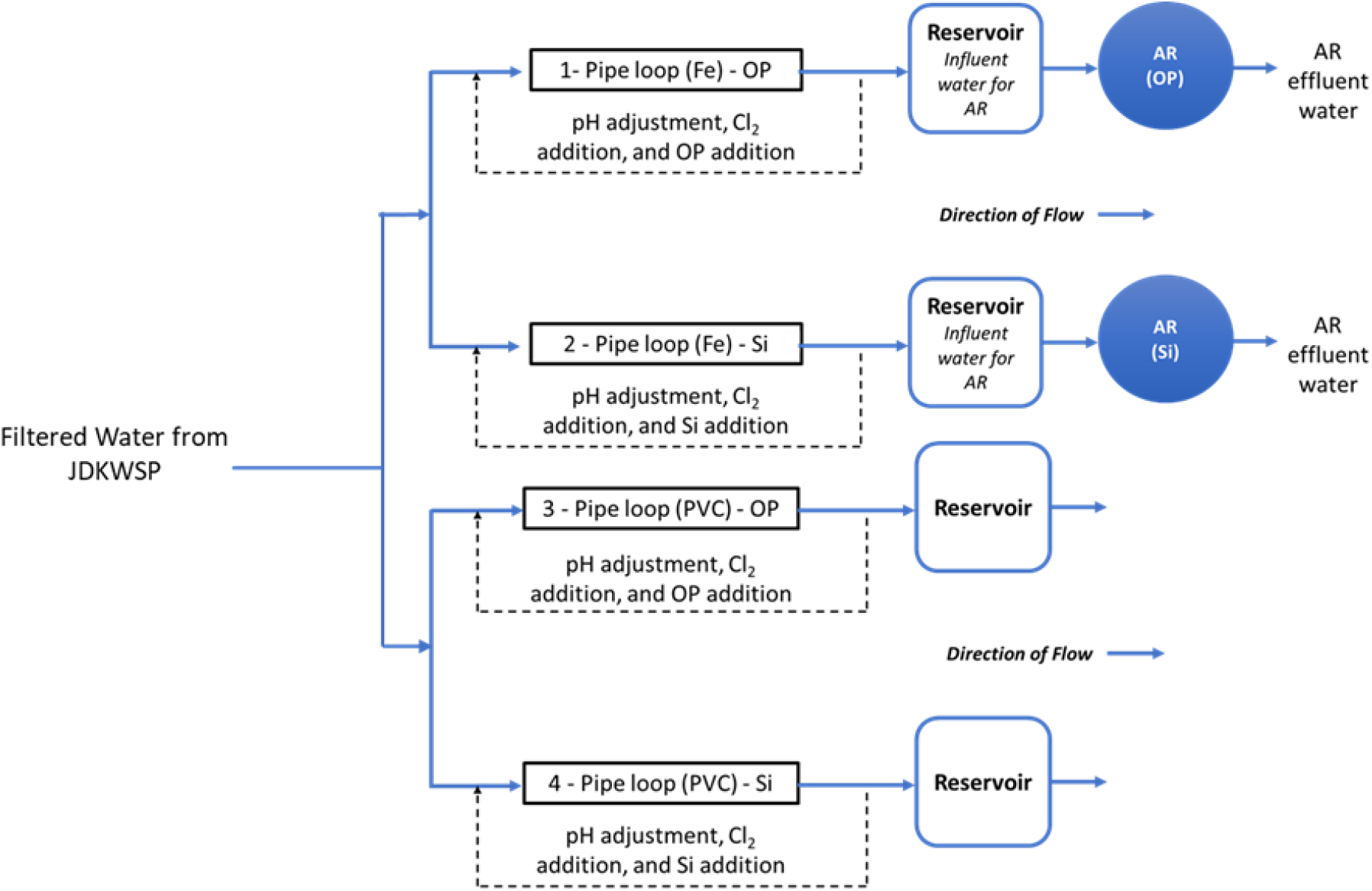
Simplified schematic of the pipe loop system.

Filtered water from JDKWSP was dosed twice per week with sodium hypochlorite, sodium hydroxide, and phosphoric acid (pipe loops 1 and 3) or sodium silicate (pipe loops 2 and 4) before entering the pipe loops to maintain a target residual concentration of 1.0 mg L^-1^ of free chlorine, pH of 7.4, 1.0 mg L^-1^ of PO_4_^3-^, and 24 mg L-1 of SiO_2_, respectively, in pipe loop effluent. The SiO_2_ residual target was increased to 48 mg L^-1^ in the final stage of the experiment to investigate the effect of sodium silicates on biofilm formation at higher doses. Further information about the model distribution system and a graphical water quality summary of this system can be found in Li *et al*.^14^.

Water collected at the reservoirs from the cast-iron pipe loops was used to feed two ARs, one for each corrosion inhibitor treatment (Figure 1). These ARs were used to assess the effect of each corrosion inhibitor on biofilm formation in a cast-iron main distribution system. Polycarbonate coupons were suspended in the reservoirs from of all pipe loops to identify the impact of substrate material on biofilm development.

#### Annular Reactors

ARs have been successfully used for bench-scale models to mimic water distribution systems and to simulate biofilm growth in a variety of aquatic environments including drinking water systems, hulls of ships, and river ecosystems.^24–27^ The ARs used in this research were manufactured and sold by BioSurface Technologies Corporation (Model 1320 LS, Bozeman, MT, USA). The ARs are made up of an inner, slotted polycarbonate cylinder with a stationary glass outer cylinder. Twenty removable polycarbonate slides were used to support biofilm growth. These coupons were extracted from their reactors and analyzed for biofilm ATP concentrations and DNA sequencing.

One AR was connected to the effluent of each cast-iron pipe loop (pipe loops 1 and 2), downstream of the reservoir, to assess biofilm development in the presence of either orthophosphate or sodium silicate. The ARs were connected in December 2018 and acclimated for 3 months before the first set of coupons and bulk water were sampled in March 2019. This acclimation period was selected to ensure biofilm growth in the presence of residual free chlorine for disinfection. A total operating volume of approximately 950 mL was used for all AR experiments. Water quality from each AR is summarized in Figure S1. With this working volume, the HRT of the reactors was set to two hours by pumping water at a flow rate of 7.92 mL min^-1^. This HRT has been commonly used in various experiments to test biofilm growth under different water treatment and operational conditions.^24,27–29^ The rotational speed of the reactors was set to 50 RPM.

Before use, the ARs, fittings, tubing, flow breaks, and coupons were thoroughly cleaned and sterilized. These components were cleaned with phosphorus-free detergent and rinsed with deionized water. All non-metal components of the ARs were soaked in 10% nitric acid solution for 24 hours and rinsed with deionized water to remove any remaining impurities. The assembled ARs with mounted coupons were fitted with Masterflex PerfectPosition Norprene tubing (Cole-Parmer Canada Company, QC, Canada) and autoclaved at 121°C for 15 minutes. Once autoclaved and cooled, all non-opaque surfaces were covered with aluminum foil to reduce phototrophic growth within the reactors.^27^ To introduce influent water into the reactors, peristaltic pumps (Cole-Parmer Canada Company, QC, Canada) were calibrated to feed the appropriate flow rates. After set-up, the reactor systems were fine-tuned to their corresponding rotational speeds and left to complete their acclimation and testing period.

One coupon from each AR was sampled every second week from March 2019 until the end of June 2019, after a three-month acclimation period that started in December 2018. The sampling frequency increased to one coupon per week after July 2019. During this period the coupons were extracted to quantify and compare total ATP (tATP) by sampling approximately 4.2 cm^2^ of each of the coupon’s exposed surface area.

The coupons were aseptically removed from the ARs with a flamed sterilized metal hook and placed in 150-mL sterile test tubes. Each test tube was filled with effluent water to prevent biofilm from drying. The test tubes carrying the coupons were transported in an ice-packed cooler to Dalhousie University for analysis. All microbiological tests were performed within 24 hours of sample collection inside a biological safety cabinet (BSC) to limit biological contamination of samples and maintain sterility of materials. Once biofilm collection from the coupons was completed, the coupons were cleaned with phosphorus-free detergent, rinsed with deionized water (Reference A+, Milli-Q, Millipore), and disinfected with 70% ethanol. The coupons were then autoclaved at 121°C for 15 minutes. Once cooled, the coupons were re-inserted into their corresponding ARs.

#### Reservoir-suspended coupons

To evaluate biomass accumulation under low water temperature conditions and different pipe materials, 14 removable polycarbonate coupons were suspended from the top of each reservoir (Figure 1). All polycarbonate coupons were autoclaved at 121°C for 15 minutes prior to being used for this experiment. The coupons were suspended and submerged in October 2019 inside the tanks using nylon fishing lines and wood clamps. The majority of the surface area of these coupons (approximately 25 cm^2^) were used to assess the biomass accumulated from each treated pipe loop system.

For this specific assessment, two polycarbonate coupons per reservoir were assessed weekly from October to November 2019 to compare the impact of pipe loop material (cast-iron vs PVC) on biomass accumulation using tATP quantification. Coupons completed a 15-day conditioning period before the first set of samples was collected and analyzed. The collection process used for the coupons from the ARs was again applied for this part of the study.

### Batch Reactors

Biofilm growth at bench-scale was evaluated using amber glass reactors with annular PEX coupons by measuring the accumulated biomass as tATP. All components of the reactors were soaked in 10% nitric acid solution for 24 hours and rinsed with deionized water. The assembled batch reactors and coupons were autoclaved at 121°C for 15 minutes. The reactors were stored in the dark and amber glass was used to limit the effects of light on phototrophic bacteria growth. Four coupons in total were inserted per reactor and each coupon had an internal surface area of approximately 28.3 cm^2^ to 30.2 cm^2^, which was used for biofilm sampling (Figure S2). The test solutions (300 mL per reactor) comprised filtered water from the JD Kline water supply plant modified with orthophosphate (2 mg P L^-1^) and sodium silicate (25 mg SiO_2_ L^-1^) at 20°C. As a final step, pH was adjusted to 8 with 0.1 M NaOH. Free chlorine was dosed at 1 mg L^-1^ free chlorine and allowed to deplete over the experiment. The solution in the reactors was replaced daily using a dump and fill procedure. The coupons were acclimated for one month, under the experimental conditions, before collecting the samples.

### Adenosine Triphosphate Analysis

Total ATP (tATP) was used as an indicator of total sessile biomass concentration accumulated on the surface of the coupons. Gora *et al*.^30^ reported that swabbing was superior to scraping for biofilm recovery; thus, the exposed surface area of the coupons was swabbed for biofilm recovery and total ATP analysis. The extracted biofilm samples were then processed using LuminUltra Technologies’ Deposit & Surface Analysis (DSA) test kit and protocol (New Brunswick, Canada), and analysed with their PhotonMaster luminometer to obtain relative light units (RLUs). RLU were then translated to tATP concentrations (pg ATP cm^-2^) using LuminUltra’s conversion formula. The detection limit for this method is 10 RLUs.

### Bioinformatic analysis

Analysis of 16s rRNA sequencing and bacterial species identification was performed on biofilm extracted from the surface of the AR coupons at the beginning of each month from June – December 2019. Seven coupons (one coupon per month, per AR) were extracted from each AR. These coupons were sampled on the first week of each month to identify any variations in the microbial community structure at the highest average tATP concentrations (July to September). For this analysis the full surface area of the coupons was swabbed to extract the largest amount of biomass. DNA extraction was performed using the procedure outlined by the commercially available DNeasy PowerBiofilm Kit (QIAGEN, Hilden, Germany). All extracted DNA samples were sent to the Integrated Microbiome Resource (IMR) Laboratory at Dalhousie University for marker gene sequencing, using PCR V4-V5 primers that target the 16s ribosomal RNA (rRNA) gene in bacteria.

Results from the microbial community sequence were processed using the Microbiome Helper workflow^31^ specific to 16S analysis obtained from GitHub (https://github.com/LangilleLab/microbiome_helper/wiki/Amplicon-SOP-v2). First, cutadapt^32^ was used to remove primer sequences from sequencing reads. The trimmed primer files were imported into QIIME2 for microbiome analysis.^33^ Then, forward and reverse paired-end reads were joined using DADA2,^34^ and input into Deblur^35^ to correct reads and obtain amplicon sequence variants (ASVs). ASVs with frequency lower than 0.1% of the mean sample depth were excluded from further analysis. MAFFT^36^ was used to build a multiple-sequence alignment of ASVs and taxonomy was assigned to ASVs using the SILVA rRNA gene database^37^ and the “feature-classifier” option in QIIME2. Further data processing and visualization was completed with the Phyloseq package in R.^38^ Alpha diversity metrics including richness, Shannon’s diversity index, and Pielou’s eveness index were also analyzed. The Shannon diversity index describes species diversity within a community while taking richness (number of species present) into account. The Pielou index ranges from 0 to 1 and indicates how even the distribution of species is within a community. When the Pileou index approaches 0, it means that the community is dominated by a small subset of the total number of species (less evenness).

### Water quality parameters

All aqueous samples were collected in 500-mL high-density polyethylene (HDPE) bottles once per week. Before use, sampling bottles were soaked in a 10% nitric acid solution for 24 hours and rinsed three times with deionized water. Samples collected from the JDKWSP were stored in an ice-packed cooler and transported to Dalhousie University (Halifax, Nova Scotia, Canada) for further analysis.

pH and temperature were measured immediately after sampling. The pH was measured using an Accumet XL-50 dual channel meter (Fisher Scientific, MA, USA) according to the manufacturer’s instructions with a calibration completed each day before use. Temperature was measured using an alcohol thermometer.

Free chlorine, phosphate, and silica were also measured immediately after sampling using a DR6000 spectrophotometer (HACH, CO, USA), following the USEPA DPD method (Method 8021), PhosVer 3 method (Method 8048), and Silicomolybdate method (Method 8021), respectively. The analytical ranges for these methods are between 0.02 to 2.00 mg L^-1^ for Cl_2_, 0.02 to 2.50 mg L^-1^ for orthophosphate, and 1 to 100 mg L^-1^ for SiO_2_.

To measure ATP concentrations in bulk water, a minimum of 50 mL of water was collected in sterile falcon tubes. Cellular ATP (cATP) was used as an indicator of planktonic cell concentrations in aqueous samples. cATP was measured using LuminUltra Technologies’ (New Brunswick, Canada) Quench-Gone Aqueous (QGA) test kit and protocol, along with their PhotonMaster luminometer. The RLUs measured were translated to ATP concentrations (pg ATP mL^-1^) using LuminUltra’s conversion formula. The detection limit for this method is 10 RLUs.

### Standards and reagents

Stock solutions of sodium hydroxide (1 M) (Fisher Scientific, MA, USA), sodium hypochlorite (Atlantic Chemical and Aquatics Inc., NB, Canada), phosphoric acid (0.05 M) (Fisher Scientific, MA, USA), and sodium silicate solution (Na_2_O:SiO_2_ = 1:3.22, weight ratio) (National Silicates, ON, Canada), were made each week and diluted with feedwater to achieve the desired residual water quality concentrations.

### Data analysis

Experimental data were analyzed using R (Version 3.6.2) (R Core Team, Vienna, Austria) and a variety of widely-used contributed packages.^38–41^ Wilcoxon-signed rank tests (wilcox.test() in R) were used to compare paired samples under the assumption that repeated measurements from individual annular reactors were independent. Violation of this assumption tends to inflate the type I error rate (incorrect rejection of the null hypothesis). Since no statistical test on AR time series data resulted in a rejection of the null hypothesis, we did not evaluate the validity of the independence assumption in detail.^42^ Correlations were calculated using Pearson’s product moment correlation coefficient (cor() in R). All statistical analyses were conducted at the 95% confidence level (*α* = 0.05).

Differences in alpha diversity (species richness, diversity, and evenness) between groups were calculated using Wilcoxon-signed rank tests with correction for false discovery rate (FDR). Beta diversity (community composition) was assessed using weighted and unweighted UniFrac principal coordinate analysis (PCoA) to explore similarities between samples. Unweighted UniFrac is based on the presence/absence of different taxa and abundance is not considered, while the weighted UniFrac considers the abundance of different taxa.^43,44^ Differences in beta diversity were calculated applying a permutational ANOVA (PERMANOVA) Adonis test, which is a non-parametric multivariate analysis of variance that compares the abundance of each taxon in a sample to its abundance in other samples.^45^

## Results and discussion

### Annular Reactors: Impact of sodium silicate on biofilm formation in a pilot-scale set-up

#### Cellular ATP (cATP) and biofilm ATP (tATP)

While cellular ATP concentrations were similar in reactors treated with sodium silicate and orthophosphate, biofilm ATP concentrations were greater in the presence of orthophosphate. Figure 2a and Figure 2b show the cellular ATP in the influent and effluent of ARs treated with either orthophosphate or sodium silicate. Cellular ATP concentrations increased in the effluent of both ARs during the July - September quarter. The median cellular ATP concentrations were 2.12 pg cATP mL^-1^ (range: 0.06 - 36.3 pg cATP mL^-1^, *n* = 66) and 1.84 pg cATP mL^-1^ (range: 0.07 - 68.0 pg cATP mL^-1^, *n* = 66) in effluent from the orthophosphate- and sodium silicate-treated ARs, respectively. A Wilcoxon-signed rank test confirmed that this difference was not statistically significant (*p* > 0.05). However, an immediate spike in effluent cellular ATP from the sodium silicate-treated AR was evident after the sodium silicate dose was doubled at the beginning of September (Figure 2). This represents the maximum recorded cellular ATP concentration in all of the systems tested (68.0 pg cATP mL^-1^); increasing the sodium silicate dose appeared to promote the dispersion and release of biofilm-bound cells from the AR into the water.

**Figure 2:**
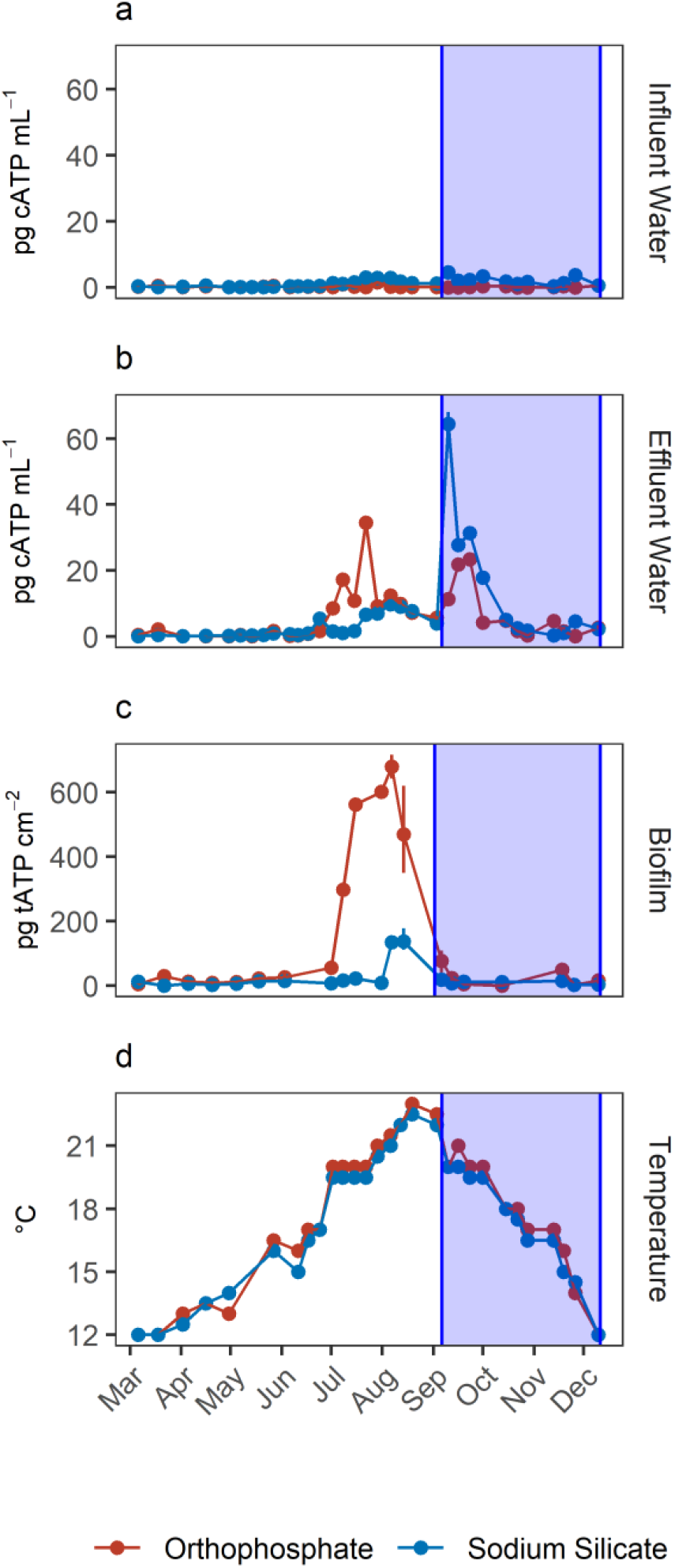
Average influent and effluent water ATP concentrations in pg cATP mL^-1^ (a and b) and average biofilm ATP concentrations in pg tATP cm^-2^ (c) from the orthophosphate- and sodium silicate-treated ARs; Error bars represent maximum and minimum values, n = 2. The shaded area represents an increase in sodium silicate dose from 24 to 48 mg SiO_2_ L^-1^.

Total ATP concentrations, which represent the total sessile biomass accumulated on the polycarbonate coupons, increased in both systems during the July - September quarter (Figure 2c). The maximum biofilm ATP concentrations values of biofilm ATP concentrations were recorded during the same quarter. Median concentrations were 30.4 pg tATP cm^-2^ (range: 0 - 717.0 pg tATP cm^-2^, *n* = 46), and 10.6 pg tATP cm^-2^ (range: 0 - 178.0 pg tATP cm^-2^, *n* = 46) in biofilm samples recovered from the orthophosphate- and sodium silicate-treated reactors, respectively. Peak biofilm ATP concentrations from the silicate-treated AR were approximately 4 times lower than the corresponding peak representing the orthophosphate-treated AR.

The increase in biomass concentrations in the orthophosphate-treated system is consistent with previous drinking water studies reporting an increase in microbial growth due to orthophosphate.^6–8,10,46,47^ Lehtola *et al*.^7^ reported an increase in biomass concentrations after adding orthophosphate as little as 1 µg P-PO_4_^3-^ L^-1^ to treated drinking water with no disinfectant residual and a background of 0.19 µg L^-1^ of available phosphorus. While Rompré *et al*.^2^ and Kogo *et al*.^19^ indicated that biofilm accumulation was not influenced by corrosion inhibitor type (sodium silicate or orthophosphate), the results from this study suggest the opposite: tATP concentrations were significantly lower in the silicate-treated AR compared to the orthophosphate-treated AR. A similar finding was documented by Aghasadeghi *et al*.^48^, who used a pilot-scale system with excavated lead service lines. In that study, heterotrophic plate counts were lower in pipe wall biofilm in the silicate-treated system compared with the orthophosphate-treated system.

Cellular ATP and biofilm ATP reported in this section were not strongly correlated (Orthophosphate = -0.23, Sodium Silicate = 0.07) (Figure 2). The two microbial indicators appear to be governed by different mechanisms, although there is some overlap in the orthophosphate system’s biofilm ATP and cellular ATP spikes.

### Reservoir-suspended coupons: Influence of corrosion inhibitor on biofilm growth at lower water temperatures

To monitor the biomass accumulated at low water temperatures (below 20°C), polycarbonate coupons submerged in the effluent reservoirs for each PVC and cast-iron pipe loop were used for biofilm ATP analysis. The average reservoir water temperature was 16.8°C (range: 16.5 -17.5 °C) and 17.3 (range: 17.0 - 18.0 °C) in the sodium silicate- and orthophosphate-treated systems, respectively. Coupons removed from the sodium silicate-treated systems had lower median ATP concentrations than those removed from the orthophosphate-treated systems regardless of the water main composition (cast-iron or PVC) (Figure 3). The median biofilm ATP concentrations representing the cast-iron pipe loop systems treated with sodium silicate and orthophosphate were 20.9 pg tATP cm^-2^ (range: 9.62 - 55.2 pg tATP cm^-2^, *n* = 12) and 31.8 pg tATP cm^-2^ (range: 0.00 - 77.8 pg tATP cm^-2^, *n* = 12), respectively. Similarly, the median biofilm ATP concentrations representing the PVC pipe loop systems treated with sodium silicate and orthophosphate were 18.2 pg tATP cm^-2^ (range: 0.00 - 46.9 pg tATP cm^-2^, *n* = 12) and 22.3 pg tATP cm^-2^ (range: 9.14 - 79.5 pg tATP cm^-2^, *n* = 12). However, a two-way ANOVA test with Tukey’s HSD method conducted on the data set indicated that there were no significant differences in average biofilm ATP concentrations between the pipe materials and the corrosion inhibitor treatments (PVC - cast-iron CI: -14.9, 5.8 pg tATP cm^-2^; Sodium Silicate - Orthophosphate CI: -16.7, 3.9 pg tATP cm^-2^). Significant differences in biofilm ATP concentrations were only evident at peak temperatures (Figure 2, July - September).

**Figure 3:**
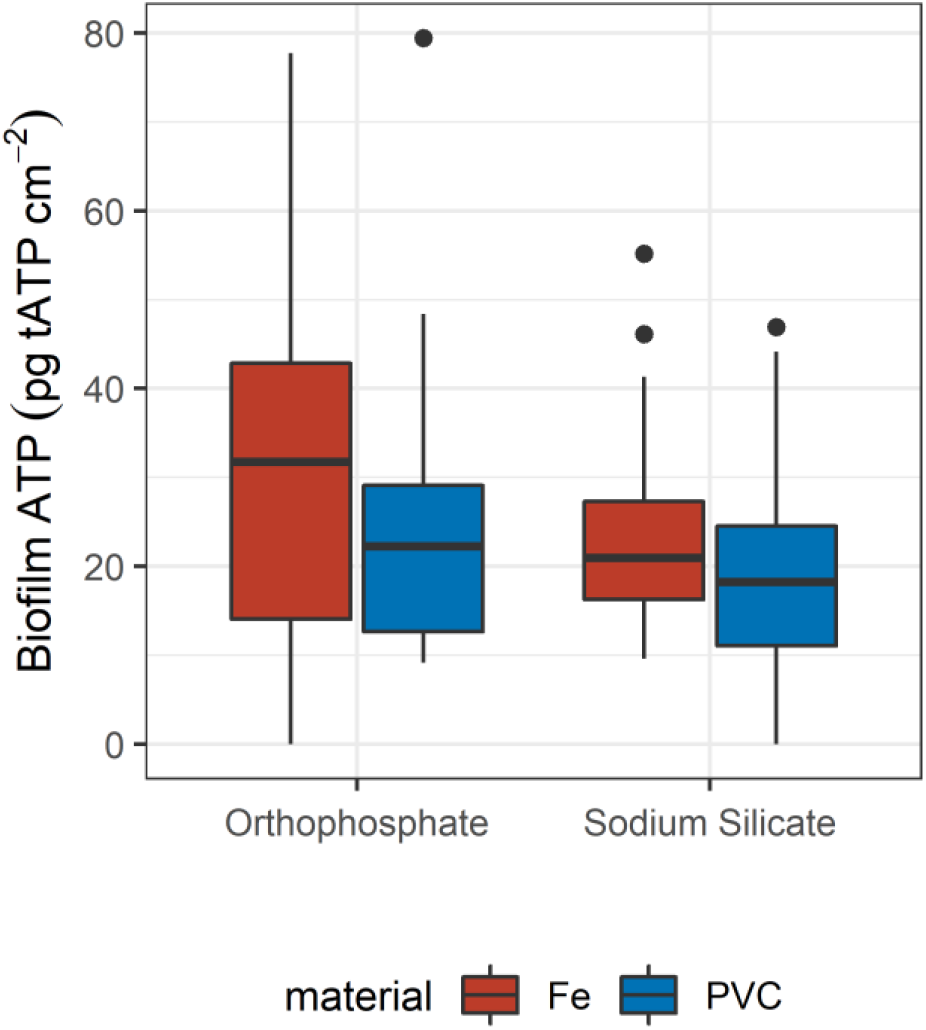
Box plot of tATP measured from the coupons submerged in the reservoirs of each PVC and cast-iron pipe loop; n = 12

While not evident in this study, pipe material does influence biofilm growth in drinking water systems.^49–53^ For example, Niquette *et al*.^51^ demonstrated that pipe material considerably influenced the density of fixed biomass and that PVC, along with polyethylene, supported less fixed biomass than iron and cement-based materials. A similar trend is shown in Figure 3, where the median tATP concentrations in the sodium silicate-treated coupons are lower than the orthophosphate-treated coupons in each of the pipe loop material categories (cast-iron vs PVC), and the median tATP concentrations are lower in the PVC compared to the cast-iron pipe loops.

### Batch Reactors: Influence of corrosion inhibitor on biofilm growth at high-water temperatures

The substantial difference between biofilm growth in the presence of sodium silicate and orthophosphate at water temperatures above 20°C was confirmed using batch reactors with PEX coupons. While orthophosphate was associated with increased tATP concentrations on PEX pipe coupons, sodium silicate was not (Figure 4). In the absence of disinfectant, the median biofilm ATP concentrations on the orthophosphate- and sodium silicate-treated coupons were 86.9 pg tATP cm^-2^ (range: 61.1 - 109.0 pg tATP cm^-2^, *n* = 4), and 52.2 pg tATP cm^-2^ (20.8 - 61.1 pg tATP cm^-2^), respectively.

**Figure 4:**
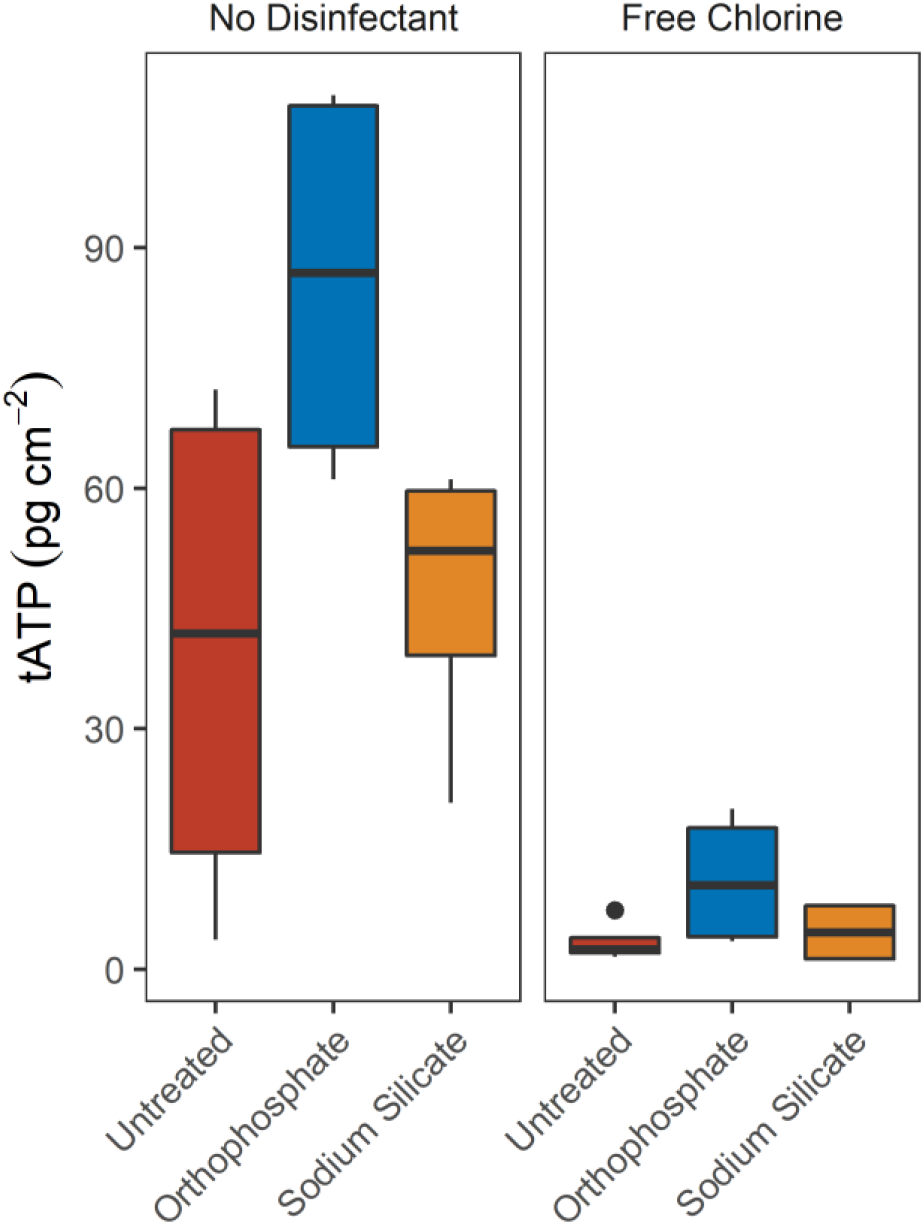
tATP concentrations on annular PEX coupons in batch reactors with modified filtered water at 20°C with and without chlorine addition; n = 4

Biofilm ATP surface concentrations were lower on coupons exposed to free chlorine, but the orthophosphate-treated coupons recorded a higher median ATP values than the untreated and the sodium silicate-treated coupons (Figure 4). These results complement the findings from the pilot-scale study in which higher biomass concentrations were recorded from the orthophosphate-treated AR when water temperature increased above 20°C during Q2 (July - September). A two-way ANOVA test with Tukey’s HSD comparison confirmed that the average biofilm ATP concentrations were significantly different between the coupons that were treated with orthophosphate and the ones that did not have any corrosion control treatment (Orthophosphate - Untreated CI: 2.02 - 51.6 pg tATP cm^-2^).

### Microbial community structure

#### Taxonomic analysis

Manganese oxidizing bacteria (MOB) were more abundant in the sodium silicate-treated system compared to the orthophosphate-treated system. The genera *Hyphomicrobium* and *Sphingomonas*, which consist of manganese oxidizing bacteria (MOB),^54,55^ were present in both orthophosphate- and sodium silicate-treated systems (Figure 5). In the case of *Hyphomicrobium*, the sodium silicate samples had a relative abundance of 4.8% and 2.8% in June and July, respectively. In comparison, the highest abundance of *Hyphomicrobium* in the orthophosphate-treated system was 0.9% in December. *Hyphomicrobium* was previously identified in biofilms collected in ARs that were operating with untreated water from Pockwock Lake (raw water from JD Kline water supply plant);^56^ whereas the most abundant MOB identified by Allward *et al*.^56^ (*Candidatus Koribacter*) was not identified in this study for any sample collected.

**Figure 5:**
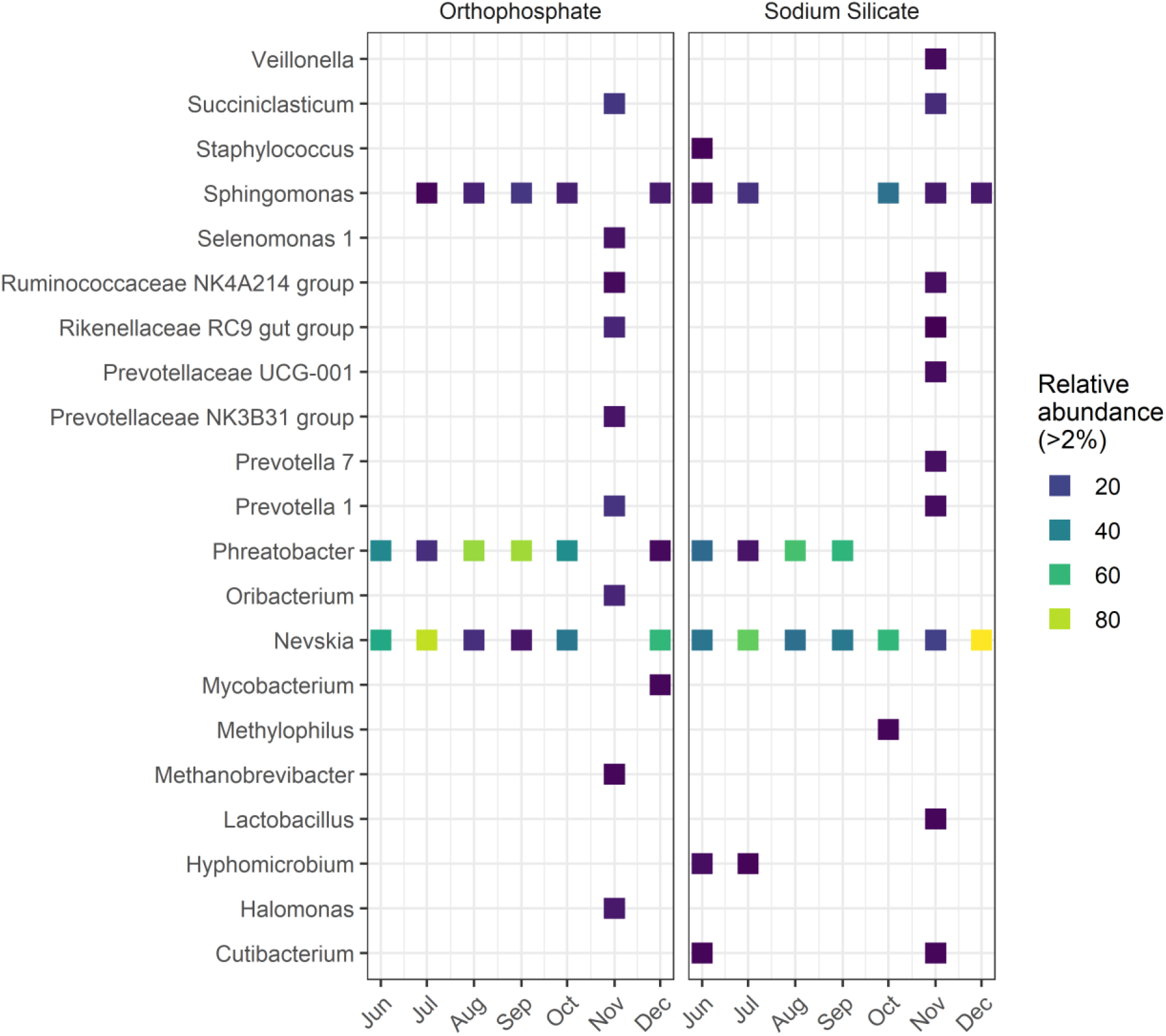
Relative abundance of bacterial genera, by month.

Similarly, the highest abundance of *Sphingomonas* was detected in samples from the silicate-treated AR collected in the month of October (34.3%). In comparison, the highest relative abundance for this genus in the orthophosphate-treated AR occurred in September (15.8%). Known iron-oxidizing bacteria (FeOB) that are part of the genus *Gallionella* and *Leptobthrix*^57^ were not identified in this study. Additional taxonomic profiles at the phylum level are summarized in Figure S3.

The genera *Mycobacterium* and *Halomonas*—which include pathogenic organisms— were more abundant in the presence of orthophosphate. *Legionella*, and *Escherichia-Shigella* were identified at the genus level with low relative abundances (Table S1). The *Mycobacterium* genus consists of a large number of species that are either opportunistic pathogens or non-pathogenic organisms that can be resistant to chlorine and are frequently detected in drinking water distribution systems.^58,59^ The relative abundance of *Mycobacterium* was bellow 2% in the sodium silicate system, whereas the orthophosphate system recorded a relative abundance above 3% in December. The *Halomonas* genus, which includes species of bacteria that may display pathogenic potential in humans,^60^ was detected in the orthophosphate-treated system in each sampling month with the exception of June. In contrast, *Halomonas* were only detected in September, October, and November in the sodium silicate samples. In the month of November, the relative abundance of *Halomonas* increased to more than 7.5% and 1.9% in the orthophosphate- and sodium silicate-treated systems, respectively. The genus *Escherichia-Shigella*, which contains *Escherichia coli*, was detected, but with low frequency and lower abundance compared to *Halomonas* and *Mycobacterium*. The highest relative abundance for these genera occurred in November in the sodium silicate-treated AR at 0.6%. Furthermore, the genus *Legionella*, which includes the pathogenic species *Legionella pneumophila*, was also detected in both systems. However, while a low abundance was recorded (similar to *Mycobacterium* and *Halomonas*), the *Legionella* in the sodium silicate system was detected in September, October, and November with an increasing relative abundance from month to month. Both *Mycobacterium* and *Legionella* have similar occurrence in water distribution systems, interactions with protozoa, and tendency to form biofilms.^59,61,62^

#### Alpha and beta diversity analysis

In general, no significant differences in diversity and richness between the orthophosphate- and sodium silicate-treated groups were detected. Richness, Shannon diversity index and Pielou’s evenness index were calculated for every sequenced sample. The alpha diversity indexes are summarized in Figure S4. The observed ASVs noticeably increased in November which reflects the change in biofilm community composition in both systems. Particularly, this shift is more evident in the orthophosphate AR. Overall, biofilm samples demonstrated the highest evenness (Pielou’s index) as well as the highest diversity (Shannon Index) in November 2019. On average, the orthophosphate system displayed the largest microbial evenness (Pielou’s index: 0.50) and diversity (Shannon index: 1.61) in comparison to the sodium silicate-treated system (Pielou’s index: 0.44; Shannon index: 1.55).

No significant difference in community structure between the orthophosphate- and the sodium silicate-treated groups was identified using the weighted (*p* = 0.75) and unweighted (*p* = 0.65) UniFrac methods (Figure S5).

## Conclusion

There are existing knowledge gaps with respect to the use of sodium silicates for corrosion control and their impacts on biomass accumulation in drinking water distribution systems. The objective of this work was to compare sodium silicate with orthophosphate for biofilm formation at constant pH. The results from this work suggest the following:

1. Orthophosphate promotes more biofilm growth than sodium silicate, but only at water temperatures above 20°C. As shown in the results from the pilot-study, the biofilm ATP concentrations recorded from the orthophosphate-treated AR were significantly higher than the concentrations recorded from the sodium silicate-treated AR between July and September, when the water temperature exceeded 20°C. Two separate experiments—involving (1) polycarbonate coupons exposed to treated water from the cast-iron and PVC pipe loops at lower water temperatures (below 20°C) and (2) PEX coupons in batch reactors exposed to warmer water temperatures (20°C)—corroborate that biomass accumulation in the presence of orthophosphate at high water temperatures only. At low water temperatures (< ∼15°C), there were no significant differences in average biofilm ATP concentrations regardless of the corrosion inhibitor treatment. However, the biofilm ATP concentrations recorded in the batch reactors confirmed that when the water temperature was maintained at 20°C there is a higher biomass accumulated on the orthophosphate-treated coupons.
2. A free chlorine residual had significant impact on decreasing biofilm ATP concentrations. This was particularly noticeable for the orthophosphate treated samples.
3. Effluent cATP concentrations spiked with the increase in sodium silicate doses from 24 to 48 mg SiO_2_ L^-1^. This suggests that changes in sodium silicate concentrations disturbed and dispersed the biofilm formed inside the AR. This may present a risk when silicate is used as a water treatment additive.
4. Orthophosphate promoted biofilm growth, but it did not disrupt the community structure in comparison to the biofilms exposed to silicates. The presence of genera that include opportunistic pathogens from the genus *Mycobacterium, Halomonas, Escherichia coli*, and *Legionella* were detected in both systems. The genus *Halomonas* and *Mycobacterium* were present at greater relative abundances in the orthophosphate-treated system in comparison to the sodium silicate system. Conversely, *Escherichia coli*, and *Legionella* were more abundant in sodium silicate-treated coupons. Even though these genera were detected, it was not possible to identify organisms at the species level.

## Associated content

### Supplemental information

Figures summarizing water quality data, taxonomic analysis, and diversity analysis.

## Acknowledgements

This work was funded by an NSERC CRD grant (CRDPJ 509252-17) in collaboration with National Silicates and Cold-Block Technologies; additional funding support was provided through the NSERC/Halifax Water Industrial Research Chair program (IRCPJ 349838-16). We acknowledge the technical expertise provided by Heather Daurie (Centre for Water Resources Studies), Andrew Houlihan (Halifax Water), Jessica Campbell (Halifax Water), Nicole Allward (Centre for Water Resources Studies), Amy Murdock, and James Dalton.

