## Supplementary information for "Effect of sodium silicate on drinking water biofilm development"

### Pilot-scale model distribution system

#### General water quality

Figure S1 summarizes free chlorine, pH, phosphate residual, silica residual, and TOC measured in samples collected from the influent and effluent water of each AR. Water quality results were summarized in a quarterly format to demonstrate any changes due to seasonal effects, or due to changes in the concentration of the corrosion inhibitors used in this study.

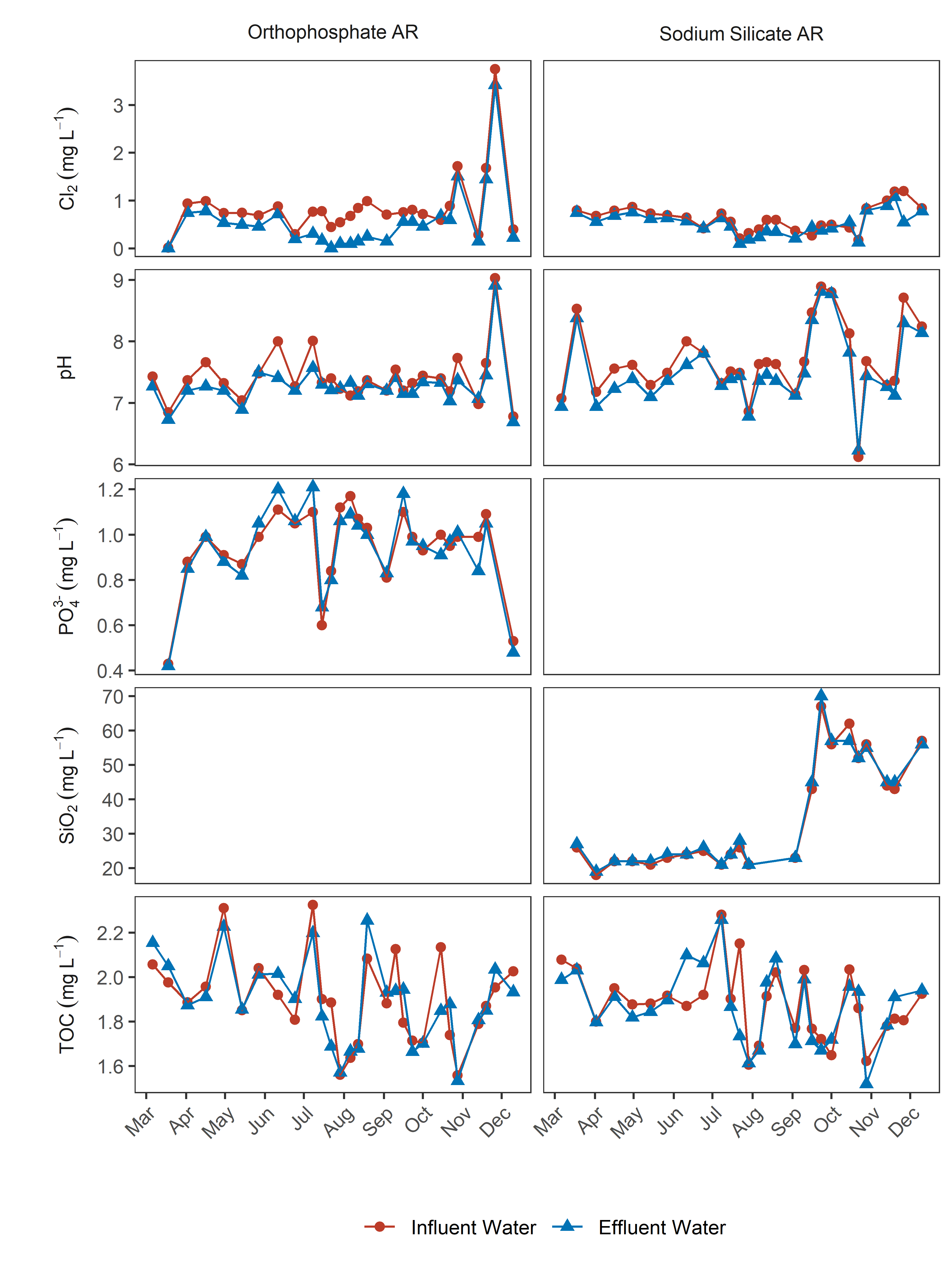

Figure S1: Free chlorine, pH, phosphate residual, silica residual, and TOC measured for the influent (reservoir) and effluent water of each AR.

**Free chlorine**

During the full extent of the study median influent free chlorine concentrations in the orthophosphate AR were 0.76 mg L^-1^ (range: 0.02 - 3.75 mg L^-1^, *n* = 26) and in the sodium silicate AR were 0.63 L^-1^ (range: 0.18 - 1.20 mg L^-1^, *n* = 26). The median influent free chlorine concentrations were significantly higher in the orthophosphate AR than in the sodium silicate reactor (*p* < 0.05). The orthophosphate-treated system also experienced a higher chlorine reduction across the AR, especially during the July - September quarter (Q2). The corresponding mean free chlorine reductions during Q2 were 69.6% in the orthophosphate AR, and 24.6% in the sodium silicate AR.

**pH**

The pH of the influent water samples gathered during the study ranged from a median of 7.35 (range: 9.03 - 6.78, *n* = 28) for the orthophosphate AR, to 7.63 (range: 8.89 - 6.12, *n* = 28) for the sodium silicate AR. The differences in pH between both systems were not significantly different at the 95% confidence level. In the sodium silicate-treated AR there was a noticeable increase in pH at the beginning of Q3 (September - December) when the sodium silicate dose increased in the pipe loop systems (September 05, 2019). This effect is explained by the alkaline nature of this type of corrosion inhibitor.

**Phosphate and silica residual**

The median phosphate residual concentrations in the influent water were stable through each quarter in contrast with the sodium silicate-treated system, mainly due to the adjustment to a higher silica residual target from 24 to 48 mg L^-1^ in September 05, 2019. The median phosphate residual concentrations were 0.99 mg L^-1^ (range: 1.17 - 0.43 mg L^-1^, *n* = 25), and the silica residual concentrations were 25.5 mg L^-1^ (range: 67.00 - 18.00 mg L^-1^, *n* = 22).

**Temperature**

Influent water temperatures recorded for each reactor system demonstrated a seasonal behavior, as expected. Median water temperatures were higher during Q2 (July - September) within each system, reaching median temperatures of 20.5 and 20.0 ^o^C for the orthophosphate and sodium silicate-treated systems, respectively. The median influent water temperatures during this study were 18.0 ^o^C (range: 23.00 - 12.00 ^o^C, *n* = 29) at the orthophosphate AR, and 17.5 ^o^C (range: 22.50 - 12.00 ^o^C, *n* = 29).

### Batch Reactors

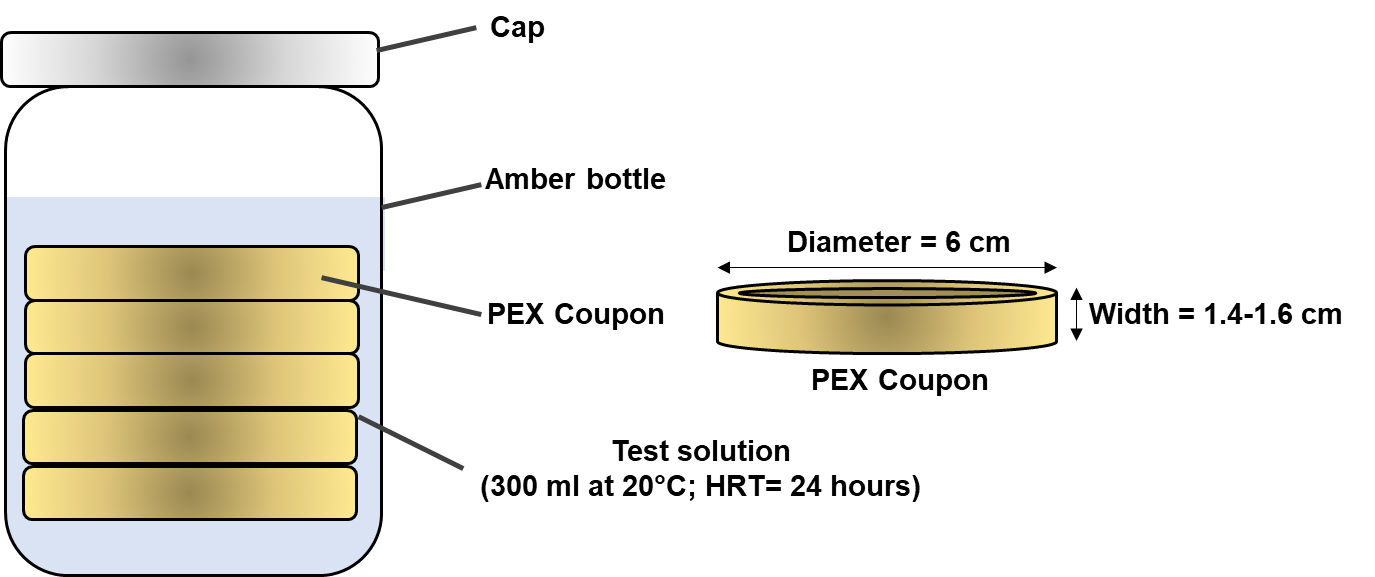

Figure S2: Simplified schematic of the batch reactors.

### Microbial community structure

#### Taxonomic analysis

The mayor ASVs identified at the phylum level were *Proteobacteria*, *Bacteroidetes*, *Firmicutes*, *Actinobacteria*, *Cyanobacteria*, and *Euryarchaeota* (Figure S2). At this phylum level, the presence of *Proteobacteria* dominated the community structure in both orthophosphate and sodium silicate-treated systems ( > 74% for every month expect for November). The presence of *Proteobacteria* was dominant from June to October until a shift in the bacterial community structure occurred in November 2019, allowing the identification of *Bacteroidetes*, *Firmicutes*, *Actinobacteria*, *Cyanobacteria*, and *Euryarchaeota*. A similar presence of organisms at the phylum level have been reported in drinking water biofilm studies using a variety of substrate materials, including ductile cast-iron, stainless steel, tuberculated cast-iron, and PVC.^1–5^ Generally, *Proteobacteria*, *Firmicutes*, *Actinobacteria*, and *Bacteroidetes* have been associated with the culturable portion of phosphate treated water on ductile cast-iron and stainless steel coupons;^2^ the same phyla were detected on both orthophosphate and sodium silicates treated cast-iron systems, however their relative abundance (with the expection of *Proteobacteria*) was higher in the orthophosphate-treated system.

The most noticeable difference at the phylum level between both orthophosphate and sodium silicate-treated systems is the presence of *Cyanobacteria* from the sodium silicate system in November, and the detection of *Euryarchaeota* from the orthophosphate system, aslo in November. Douterelo *et al.*^1^, reported that *Cyanobacteria* were positively correlated with TOC levels and was present in plastic pipes during the winter months (low water temperatures), which may suggest that with an absence of phosphorus in the sodium silicate-treated system *Cyanobacteria* was able to assimilate available carbon more effectively.

At the genus level the abundance of *Phreatobacter* was higher in the orthophosphate system (identified from June through December, except for November), than in the sodium silicate system (Figure 5). According to Perrin *et al.*^5^, the presence of *Phreatobacter* in drinking water distribution systems is related to warm water temperatures (>15 ^o^C).

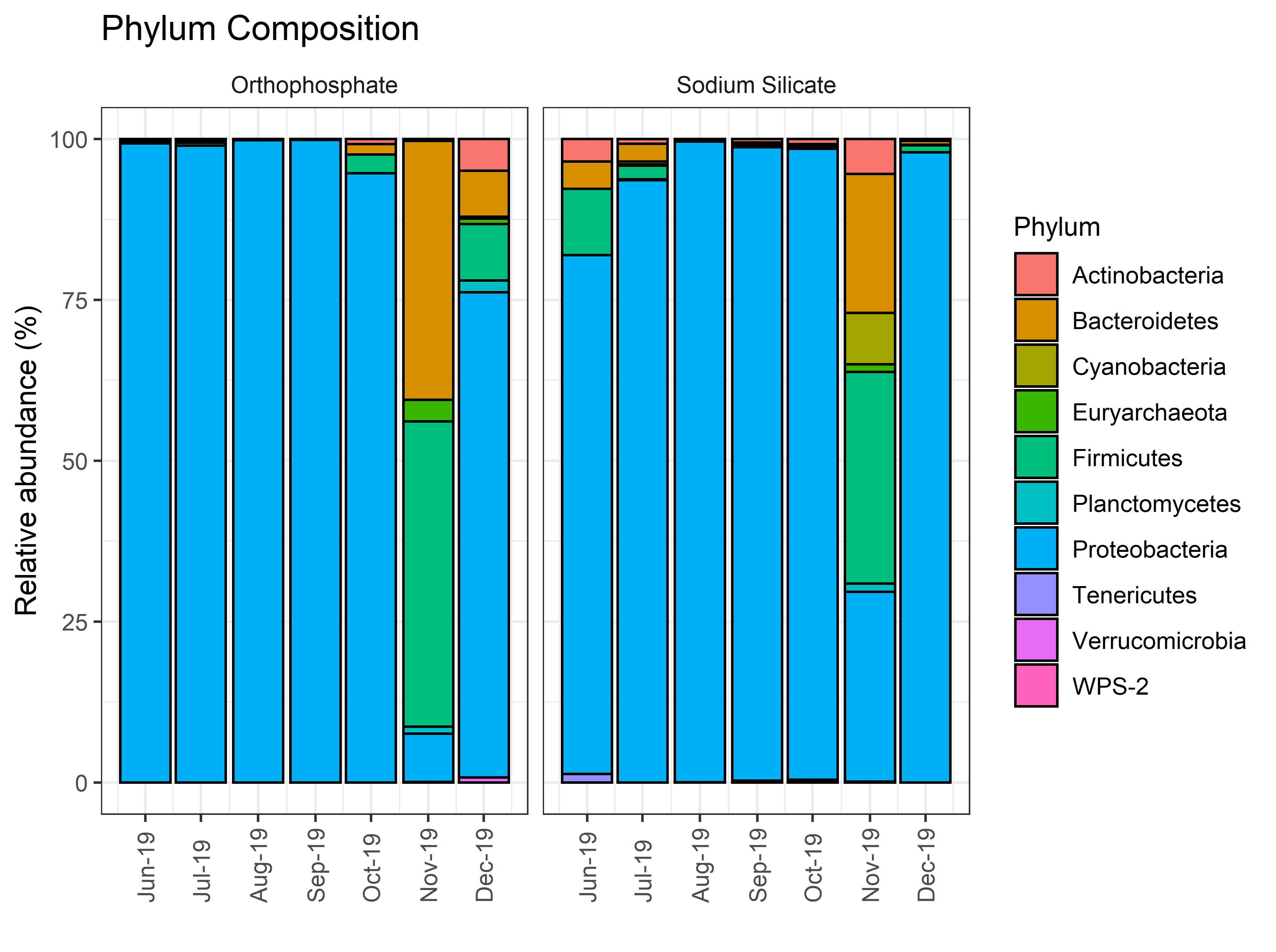

Figure S3: Relative abundance at the phylum level of bacterial community structure.

Table S1: Relative abundance (%) summary of genera associated with MOB and pathogenic bacteria species identified from the sodium silicate and orthophosphate-treated AR coupons during the months of June - December 2019.

| Month | Corrosion Inhibitor | Escherichia-Shigella | Halomonas | Hyphomicrobium | Legionella | Mycobacterium | Sphingomonas |
| --- | --- | --- | --- | --- | --- | --- | --- |
| June | Orthophosphate | - | - | - | - | - | 1.6 |
| July | Orthophosphate | - | 0.1 | 0.2 | - | 0.2 | 3.2 |
| August | Orthophosphate | - | 0.0 | 0.0 | - | 0.0 | 10.2 |
| September | Orthophosphate | 0.0 | - | 0.0 | 0.0 | 0.0 | 15.8 |
| October | Orthophosphate | 0.2 | 0.6 | 0.2 | - | 0.4 | 9.6 |
| November | Orthophosphate | - | 7.6 | - | - | - | - |
| December | Orthophosphate | 0.1 | 0.5 | 0.9 | 0.1 | 3.3 | 7.9 |
| June | Sodium Silicate | 0.2 | - | 4.8 | - | 0.3 | 5.7 |
| July | Sodium Silicate | - | - | 2.8 | - | 0.6 | 14.4 |
| August | Sodium Silicate | - | - | 0.3 | - | 0.0 | 2.0 |
| September | Sodium Silicate | - | 0.0 | 0.2 | 0.0 | 0.0 | 0.9 |
| October | Sodium Silicate | - | 0.0 | 0.2 | 0.2 | 0.1 | 34.3 |
| November | Sodium Silicate | 0.6 | 1.9 | - | 0.2 | 1.7 | 7.8 |
| December | Sodium Silicate | - | 0.0 | 0.3 | - | - | 8.0 |

#### Alpha and beta diversity analysis

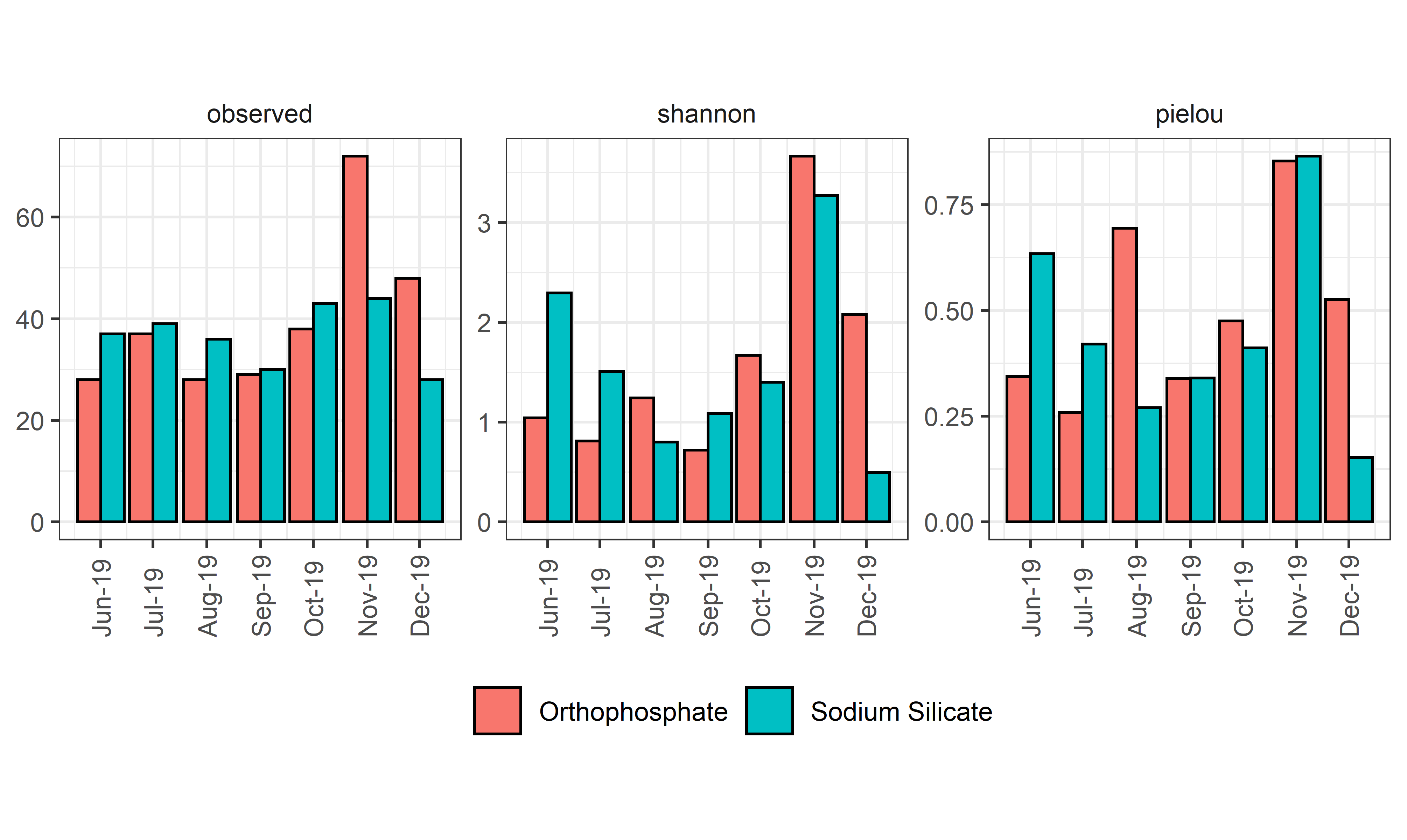

Figure S4: Richness, Shannon diversity index, and Pielou evennes index in biofilm samples collected from June to December 2019.

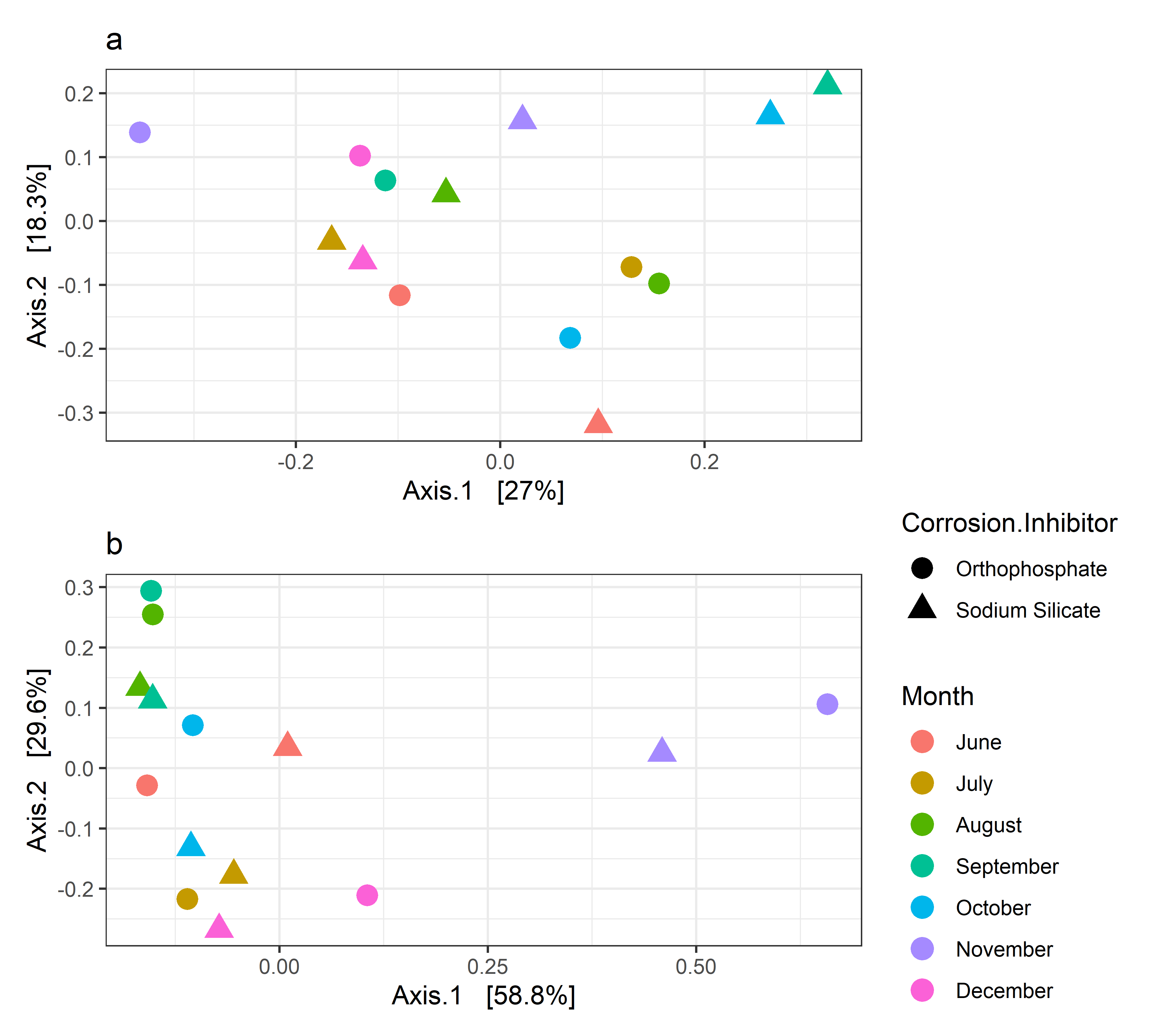

Figure S5: (a) Principle coordinate analysis for unweighted and (b) weighted UniFrac distances.
